## Supplementary figures and tables for "A spontaneous genetically-induced epiallele at a retrotransposon shapes host genome function"

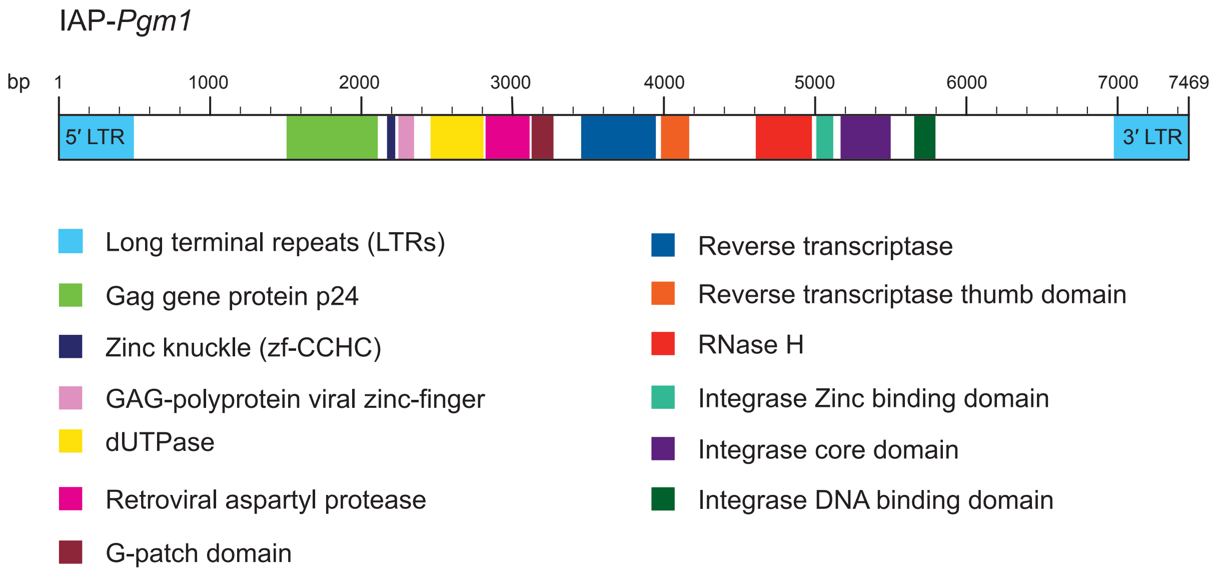


**Figure 1 - figure supplement 1.** Genomic structure and coding potential of IAP-*Pgm1*. Details lifted from the Pfam database (https://pfam.xfam.org). Genomic distances are drawn to scale.

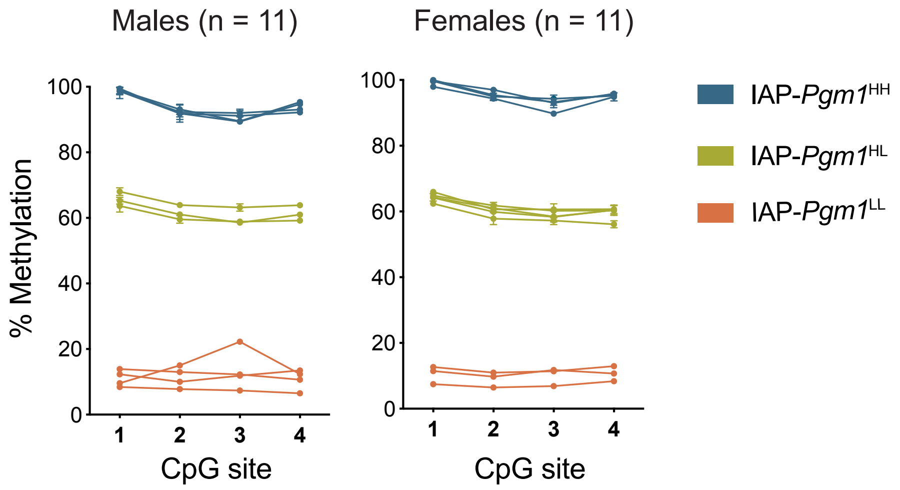


**Figure 1 - figure supplement 2.** The IAP-*Pgm1* methylation pattern is not sex-linked. IAP-*Pgm1* methylation levels in both B6 males and females segregate into three distinct DNA methylation states: high (blue), intermediate (green), and low (orange).

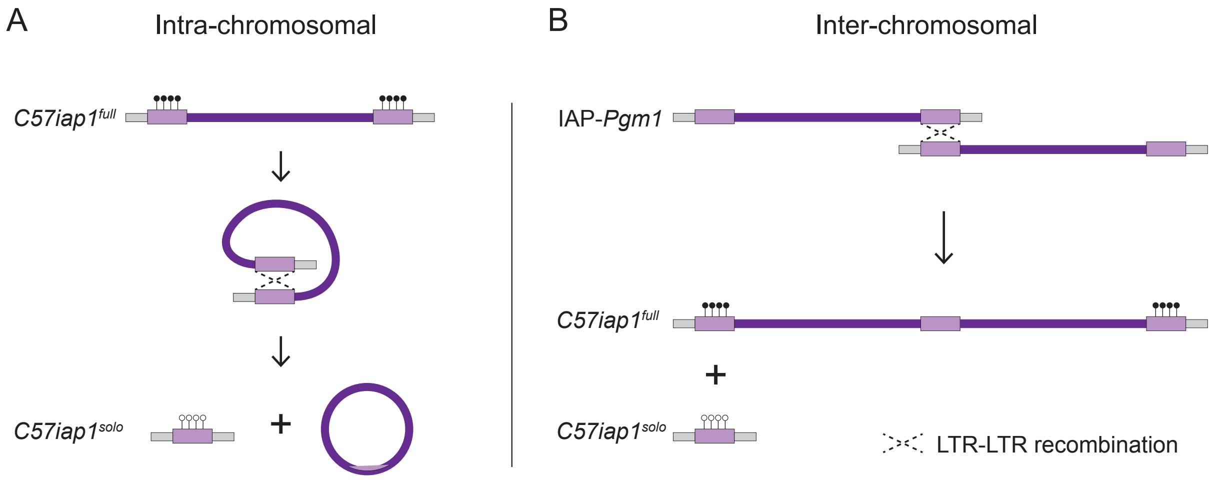


**Figure 2 – figure supplement 1.** Possible mechanisms of homologous recombination between IAP LTRs leading to the formation of a solo LTR at the *C57iap1* locus. (**A**) Intra-chromosomal homologous recombination between the 5′ and 3′ LTRs of the repeat element, giving rise to a solo LTR and a circularised unintegrated viral fragment. (**B**) Inter-chromosomal recombination between identical LTRs on sister chromatids or homologous chromosomes, producing a solo LTR on one DNA strand and a tandem duplication on the other. Diagrams adapted from Seperack et al., 2006.

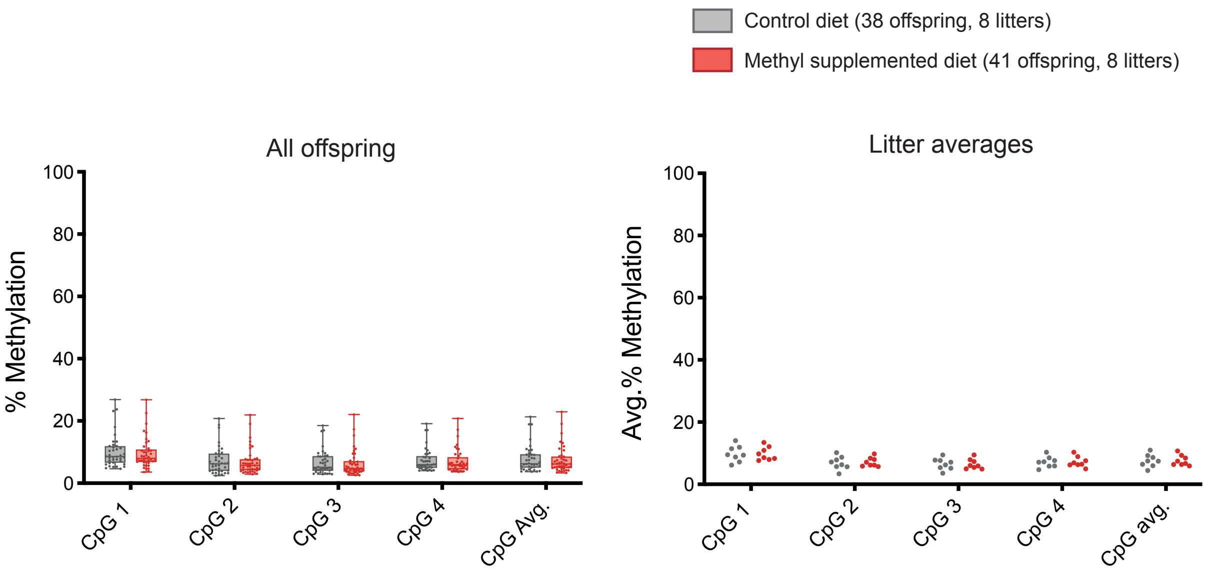


**Figure 4 – figure supplement 1.** DNA methylation at *C57iap1*$^{solo}$ is unresponsive to maternal dietary methyl supplementation. *C57iap1*$^{solo}$ females were fed a control or methyl supplemented diet two weeks prior to mating and for the duration of pregnancy and lactation. DNA methylation levels were quantified in 8-week old offspring liver samples at the four distal CpGs of the 5′ end of *C57iap1*$^{solo}$. Statistics: Unpaired t tests on litter averages for each CpG and for the CpG average.

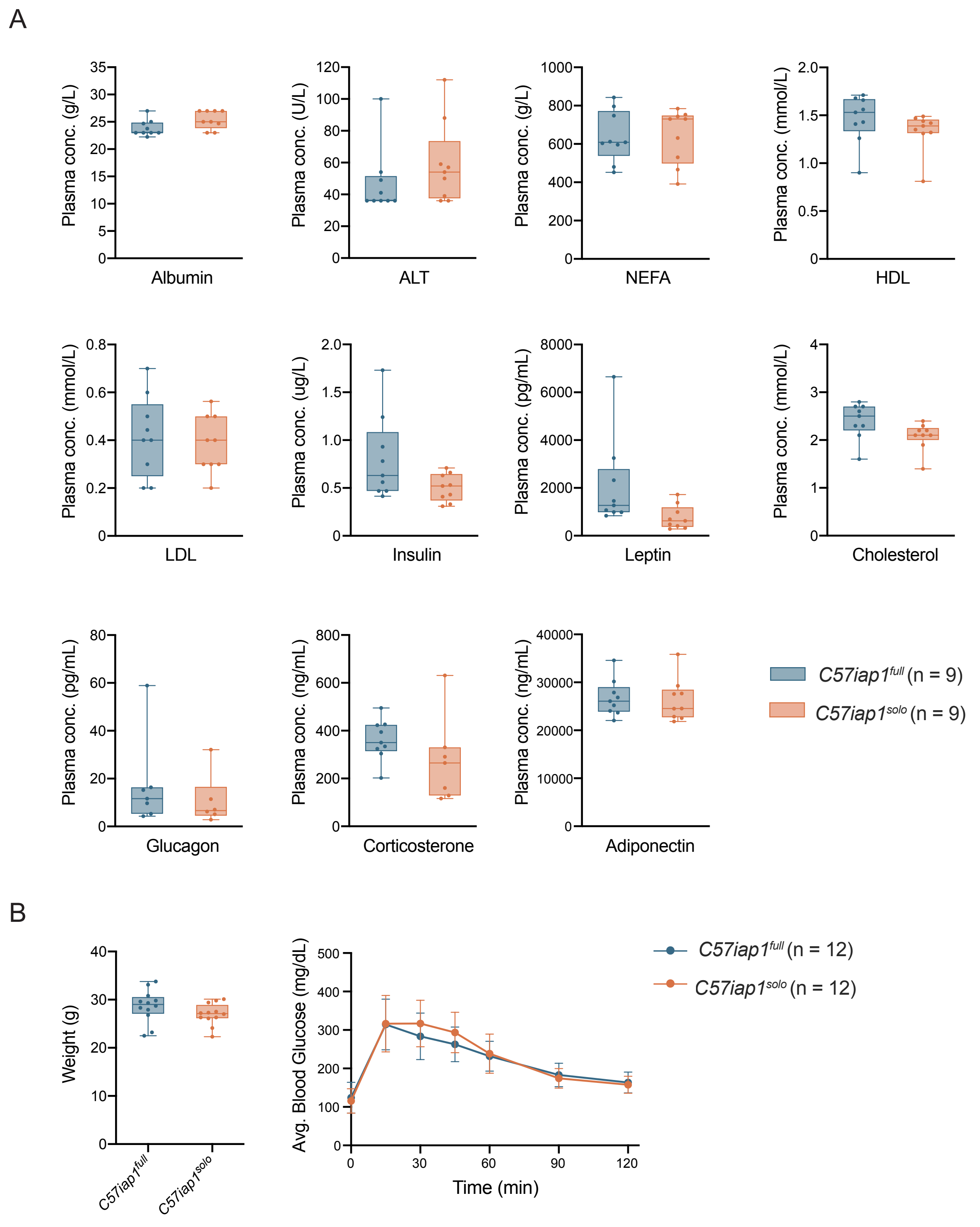


**Figure 5 – figure supplement 1.** Metabolic phenotyping of *C57iap1*$^{full}$ and *C57iap1*$^{solo}$ individuals. (**A**) Concentration of metabolic biomarkers in plasma samples collected from *C57iap1*$^{full}$ and *C57iap1*$^{solo}$ adult males. Statistics: unpaired t tests. (**B**) Glucose tolerance test on *C57iap1*$^{full}$ and *C57iap1*$^{solo}$ adult males. Following an overnight 15 hour fast, mice were weighed (left panel) and fasting tail blood glucose was measured prior to administering an intraperitoneal 2 g/kg glucose injection. Tail blood glucose measurements were taken 15, 30, 45, 60, 90, and 120 minutes from the time of injection (right panel). Statistics: unpaired t test (left panel) and multiple unpaired t tests corrected for multiple comparisons using the Holm-Šidák method (right panel).

**Supplementary Table 1. PCR primers.**

| Locus | PCR forward primer | PCR reverse primer |
|---|---|---|
| **IAP-*Pgm1* amplification** | | |
| IAP-*Pgm1*_ P1 | ACTTCTCTAACAACGTTGAGCA | CAAAGGCCTTAACTCAGCGG |
| IAP-*Pgm1*_ N1 | GAGGGGCTAGGTGGGAATAC | CCTGTATCAGAAATTTGGACC |
| IAP-*Pgm1*_ P2 | AGTCCTACAATGTGCCAACTT | GCATTCAATGTTTACCTGCCAC |
| IAP-*Pgm1*_ N2 | AGAAGCTCTTTTGCCAGTGAG | TTTCTCTTGGCGCACAGTTC |
| IAP-*Pgm1*_ P3 | ACTTCTCTAACAACGTTGAGCA | GCATTCAATGTTTACCTGCCAC |
| IAP-*Pgm1*_ N3 | GAGGGGCTAGGTGGGAATAC | TTTCTCTTGGCGCACAGTTC |
| **Genotyping** | | |
| *C57iap1$^{full}$* | TTTCATGAAGGTTCAGTGTCCT | CAACAATCCATACTCAAGTGTCC |
| *C57iap1$^{solo}$* | CAATGGCATGTGCTTAGTGG | CAACAATCCATACTCAAGTGTCC |
| Both *C57iap1* variants | CCAAGAGAAGAGAAATTACCTGGA | TCACGGGAAAGGTAGAGTAC |
| *Sry* | GCAGGCTGTAAAATGCCACT | TTCCAGGAGGCACAGAGATT |

**Supplementary Table 2. Sanger sequencing primers.**

| Primer ID | Sequence |
|---|---|
| IAP-*Pgm1* _S1 | GAGCAAGAGCCAGCTGTTTT |
| *IAP-Pgm1* _S2 | CAATGGCATGTGCTTAGTGG |
| IAP-*Pgm1* _S3 | CCAAGAGAAGAGAAATTACCTGGA |
| IAP-*Pgm1* _S4 | ACATTCGCCGTCACAAGAT |
| IAP-*Pgm1* _S5 | CTCTCTCGCTGGCATCTCTC |
| IAP-*Pgm1* _S6 | ATTGAAGGGGCTTTCGTTTT |
| IAP-*Pgm1* _S7 | CCAGTGAGTACAGCTTTACGAGGT |
| IAP-*Pgm1* _S8 | GGCCTTTCCAAGGGTCTT |
| IAP-*Pgm1* _S9 | GAAGAAGCAGCCCATTACCA |
| IAP-*Pgm1* _S10 | CTGCGTAGTGCGTCAGCAAT |
| IAP-*Pgm1* _S11 | CGTAAATACGGAACCAATGC |
| IAP-*Pgm1* _S12 | GCAGCTGCTTTGACTCCAG |
| IAP-*Pgm1* _S13 | AGCTGCGCCTCTGATAGAAC |
| IAP-*Pgm1* _S14 | GGAAAGCCTGGGCATTTTA |
| IAP-*Pgm1* _S15 | ACCCAGGAAGCAGTCAGAGA |
| IAP-*Pgm1* _S16 | GTCCTGTGCTCAAGCCCTAA |
| IAP-*Pgm1* _S17 | TCCTTGATACCGGAGCAGAT |
| IAP-*Pgm1* _S18 | AGGACAGCAAGGGAAATTCA |
| IAP-*Pgm1* _S19 | TCCAGTGTGGGTTCCTCAAT |
| IAP-*Pgm1* _S20 | GGGGTCTCCCTGTACTTTCC |
| IAP-*Pgm1* _S21 | GAGGGAACAATTCCCCTCTT |
| IAP-*Pgm1* _S22 | TGTGCCAACTTTATGTGCAAG |
| IAP-*Pgm1* _S23 | TTTTCAAAAGCTGTTGGGAGAT |
| IAP-*Pgm1* _S24 | TGGCAGCCACATCTAATGAT |
| IAP-*Pgm1* _S25 | CCTCAAGTGGTAGAATGTTTAGTGG |
| IAP-*Pgm1* _S26 | TAGAGCCCATTCAGGCCTAC |
| IAP-*Pgm1* _S27 | AAAGCTGCTGTGAGTTCTTGC |
| IAP-*Pgm1* _S28 | TCAAAAATTCCAACAGTTCTGC |
| IAP-*Pgm1* _S29 | CCAGATAGGCCCAATGAGAT |
| IAP-*Pgm1* _S30 | GTCCCTCGTCTTGGTGATGT |
| IAP-*Pgm1* _S31 | GACAGCCTTGGCTCTGTCTG |
| IAP-*Pgm1* _S32 | TGGATGGCTGAATTTGAACA |
| IAP-*Pgm1* _S33 | AATAGGTCGCTGGCCACTC |
| IAP-*Pgm1* _S34 | TTTCATGAAGGTTCAGTGTCCT |
| IAP-*Pgm1* _S35 | TAATCTGCGCATGAGCCAAG |
| IAP-*Pgm1* _S36 | TCTCTCGCTGGCATCTCTCT |
| IAP-*Pgm1* _S37 | GGACACTTGAGTATGGATTGTTG |
| IAP-*Pgm1* _S38 | CGGCAGCTATCAGAACACAA |

**Supplementary Table 3. Pyrosequencing primers.**

| Locus | PCR forward primer | PCR reverse primer | Pyrosequencing primer |
|---|---|---|---|
| IAP-*Pgm1*_ 5′ LTR | GTTTGGTTTTTATATAGAAGGAAAGAAGAA | [Btn]ACAAATAATCATAAAATACCCTTAACTCAT | ATGTTTGTTTGTTGGG |
| IAP-*Pgm1*_ 5′ border_CpG-1 | AGTAAGATAGTGGAATTATGAGAGAA | [Btn]ACATTTCTTTTCTTCTTTCCTTCTATAT | ATAAAAATGAAGATGTATAATTAGA |
| IAP-*Pgm1*_ 5′ border_CpG-2 | TTTTTATGGGTTTTGGGGATTAAATTGAAG | [Btn]CTCTAACCACACAAATACCCTAACATAAAC | AGTAAGTGTTTTTAGGGT |
| IAP-*Pgm1*_ 5′ border_CpG-3,-4 | TTATAGATTTTTTATTGTAAAGGGGGTGGT | [Btn]CTCTAACCACACAAATACCCTAACATAAAC | GGGGTGGTGTTAGTA |
| IAP-*Pgm1*_ 5′ border_CpG-5,-6,-7 | AGAGTAAAGGAGGTTTGTAGAGTAT | [Btn]ACCCCTCAAATCTACATTTACATTCC | GGTTTGTAGAGTATAGTTTAATG |
| IAP-*Pgm1*_ 5′ border_CpG-8,-9 | GGTGGAAAGTTTTTGTGTTAGGAAAATA | [Btn]CTATACTCTACAAACCTCCTTTACTC | GGAAAATAGTGGAGGTTTTA |
| IAP-*Pgm1*_ 3′ LTR | TGGTTTTTAAAGATGTAAGTAATAAAGTT | [Btn]ACAACAATCCATACTCAAATATCC | GTAGAAGATTTTGGTTTGT |
| IAP-*Pgm1*_ 3′ LTR_CpG1,2,3 | TGGGTATATTTTTAAATTGAGGTTGAGTT | [Btn]ATAACAAAAACCTAAAACAACTAATCC | AGATGGTTATAGTTATTATGTG |
| IAP-*Pgm1*_ 3′ LTR_CpG4,5 | AGGGTAAAGTATTTGATAAGAGGAAGAG | [Btn]AAACCACAAAAAAACCAACTACTT | ATTTGATAAGAGGAAGAGTT |
| IAP-*Pgm1*_ 3′ LTR_CpG6,7 | AGAAGTTGAAAAAGGTATTTAATGTTTATT | [Btn]AAAAACTCTTCCTCTTATCAAATACT | TGTTATTTTTTTAATAAATTGATAG |
| IAP-*Pgm1*_ 3′ LTR_CpG8,9,10 | GTTATTGTTTTTATTGTTGGGATAAAGTAT | [Btn]CAATCTTCTATACCAAAAAATATCTCCT | GTATAAGGTTAAGGTTGAGAA |

**Supplementary Table 4. qPCR primers.**

| Locus | PCR forward primer | PCR reverse primer |
|---|---|---|
| **RT-qPCR** | | |
| *β-actin* _exon 6 | ACGCAGCTCAGTAACAGTCC | GTGGATCAGCAAGCAGGAGT |
| *Hprt1* _exon 9 | GAGGAGTCCTGTTGATGTTGCCAG | GGCTGGCCTATAGGCTCATAGTGC |
| *Pgm1* _exon 9 | GGGATGGTGGCTCTTCACTT | CCTTCCTTCAGGGCTATGGC |
| *Rell1* _exon 7 | ATAGCCCAGCCCTCTCTTCA | CCCTTGAGACTTCCGCTGTT |
| *Tbc1d1* _exon 3 | CCTTCTCACAGCCTGGACTG | TCCATGTCTGTGGGCTGAAC |
| *5830416l19Rik* _exon 3 | CTGGAGGCGGACATTTCACT | CACCTGCTTAGCTCTGCCTT |
| **ChIP-qPCR** | | |
| *C57iap1*_ 5' end | ACAGCTGGCATGGTCTCTTC | CTCCCAACAGACAGGCATTT |
| *C57iap1*_ 3' end | GCGTGAGAACGCGTCTAATA | ATGCCCAGAGGTTTGTTTTC |
| SINE-*Rbak* _5' end | ACCAAAGTCCTGGGGTTAGC | TTGTAAGCTGCCAAACATGG |
| IAP-*Asxl3* _5' end | CACATGCGCAGATTATTTGTTT | CCAAAATACTACGCCAGTGTCA |
| *Gapdh* promoter | CATCCAGGGACGTGCTGACT | TGTGTTCTCCCCTCACTGATCTC |

**Supplementary Table 5. GEO accession numbers for ChIP-seq datasets.**

| GEO accession number | Antibody | Cells |
|---|---|---|
| GSM3173720 | HA (Gm14419-HA) | B6 ES cells |
| GSM3173728 | HA (Gm8898-HA) | B6 ES cells |
| GSM3173732 | HA (Zfp429-HA) | F9 EC cells |
| GSM3173661 | KAP1 | B6 ES cells |
| GSM3173662 | KAP1 | B6 ES cells |
| GSM3173663 | KAP1 | B6 ES cells |
